# Cytokine-driven cellular polarization in preeclampsia revealed through the “Dictionary of immune responses”

**DOI:** 10.1101/2025.03.26.645616

**Authors:** Sheng Wan, Tianfan Zhou, Yingxuan Ma, Ying Chen, Chenchen Zhou, Jing Peng, Li Lin, Weijia Luo, Wei Gu, Zhiwei Liu, Xiaolin Hua

## Abstract

**Background:** Preeclampsia (PE) is a pregnancy-related hypertensive disorder and a leading cause of maternal and perinatal mortality. Current treatments focus primarily on symptom management, as delivery remains the only definitive cure. This underscores the urgent need for innovative therapeutic strategies. Cytokines released by placental immune cells may contribute to the progression of PE and represent promising therapeutic targets.

**Methods:** We conducted single-cell sequencing on placental tissues obtained via cesarean section from patients with severe PE and cases of non-infectious preterm birth. Machine learning was applied to identify critical immune cells. Immune Response Enrichment Analysis was performed on these key cells to evaluate their polarization states and cytokine responses associated with PE. Additionally, we investigated cell–cell communications, key genes, and their related functions

**Results:** We identified cell type-specific alterations in PE, including an increased proportion of CD4+ T cells polarized towards an IL-1α- and IL-1β enriched T4-c state. Altered SPP1 signaling between macrophages and CD4+ T cells was observed, indicating immune response dysregulation. Additionally, machine learning algorithms identified four hub genes, and six small-molecule drugs were predicted to hold therapeutic potential for PE.

**Conclusions:** This study emphasizes the importance of further understanding CD4+ T cell dynamics in preeclampsia, providing potential insights into the principles of placental immunity and contributing to the exploration of immunotherapeutic strategies

## Introduction

Preeclampsia (PE) is a significant hypertensive disorder of pregnancy, affecting approximately 2-8% of pregnancies worldwide, and is associated with considerable maternal and fetal morbidity and mortality^[1,2]^. Severe preeclampsia represents a critical form of this condition, characterized by markedly elevated blood pressure and multi-organ dysfunction, which can lead to severe complications such as eclampsia, fetal growth restriction, placental abruption, and even death^[3,4]^. Its pathophysiology remains poorly understood, necessitating further investigation into its underlying molecular mechanisms and genetic regulation^[5–7]^. Current treatment options are limited primarily to the management of symptoms, as the only definitive cure is delivery, which highlights the urgent need for novel therapeutic strategies ^[8–10]^.

A commonly recognized trait in PE is an activated and modified fetomaternal immune system^[11–13]^. Understanding the immune landscape in PE not only enhances our comprehension of its etiology but also provides a foundation for developing novel therapeutic strategies aimed at modulating immune responses to improve maternal and fetal outcomes ^[13,14]^. Although the exact mechanism of PE remains unclear, the placenta plays a central role in the disease, as symptoms typically resolve following its delivery ^[15]^. Hence, maladaptation of the placental immune environment is a critical contributor to the onset and progression of PE, highlighting the urgent need for further research into the immunological aspects of the placenta . Studies have demonstrated alterations in placental immune cell populations in PE, such as elevated levels of pro-inflammatory cytokines and changes in T cell subsets ^[16,17]^. Furthermore, impaired placental implantation and abnormal uterine spiral artery remodeling lead to placental ischemia and maternal endothelial dysfunction in PE^[2,18,19]^. Placental immune cells are thought to contribute to these processes by secreting cytokines ^[20]^. Thus, the role of cytokines in the progression of PE is of significant interest. Cytokines act as crucial mediators of intercellular communication within the immune system and represent important therapeutic targets^[21,22]^. Cytokine-based therapies, including antagonists, are widely used to treat various disorders, such as cancer and autoimmune diseases^[23]^. Increasing evidence underscores the pivotal role of cytokines in shaping immune responses^[24,25]^. However, previous studies often lacked a comprehensive analysis of immune cell-specific cytokine activity across all immune cell types. To address this gap, a "dictionary of immune responses" was developed to map the global interplay between immune cells and cytokines at a single-cell resolution. This innovative resource identifies the most active cytokines in disease states and reveals how different immune cells execute distinct functions based on the cytokine signals they receive^[23]^. The emergence of this "dictionary" provides a novel perspective for investigating cytokine-immune cell polarization in disease contexts, enhancing our understanding of the dynamic roles of immune cells in disease progression.

To date, there are few studies that have depicted systematic alterations of placental immunity or investigated cytokine-immune cell polarization in PE. Through advancements in scRNA-seq methodologies, we are now capable of delineating heterogeneous biological systems with elevated precision. Some attempts have been made to ascertain the role of immune cell subsets in PE using scRNA-seq. However, most current studies in this field used normal term deliveries rather than non-infectious preterm births as the control group^[16,26,27]^, and included all types of PE instead of focusing on a specific type, such as severe or early-onset PE, which might have entirely different etiologies^[28]^. Such sample selection criteria overlook the impact of gestational age and infection factors on placental immunity, and disregards the heterogeneity of preeclampsia. The placenta sits at the interface between the maternal and fetal vascular beds and governs adverse pregnancy outcomes, encompassing PE^[29]^. while to the best of our knowledge, systematic "immune-cytokine" correlations of placenta in PE have not been well characterized by scRNA-seq.

This study aims to investigate the cytokine-driven immune cell polarization characteristics associated with PE and their related genes and signaling pathways through comprehensive analyses of single-cell and single-nuclei RNA-sequencing data. By employing various advanced multifaceted in silico approaches, immune response enrichment analysis(IREA) and cell communication analysis included, we seek to elucidate the underlying mechanisms of immune dysregulation in preeclampsia

## Materials and Methods

The flowchart depicted in Figure 1 delineates the systematic approach adopted in this research.

**Figure 1.**
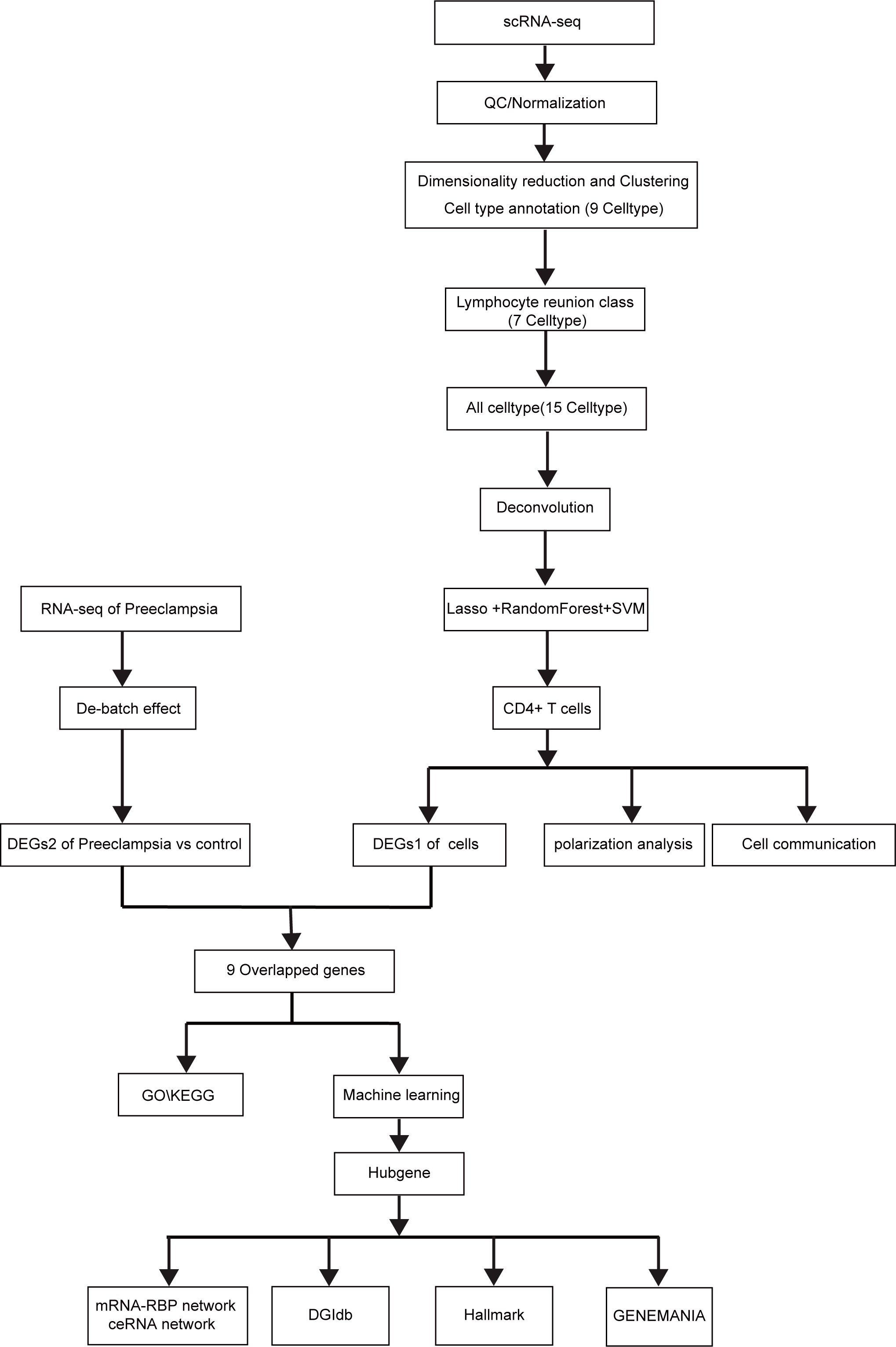
Flowchart of this study.

### Sample collection and single-cell transcriptome data acquisition

We collected six human placenta tissue samples comprising three m1 (non-infectious preterm birth) samples and three m2 (severe PE, sPE) samples in this study. Severe PE were diagnosed according to ACOG Practice Bulletin No. 222: Gestational Hypertension and Preeclampsia (2020)^[30]^. All pregnancies were delivered via Caesarean section. Detailed characteristics of the participants are provided in Table 1. The study was approved by the Ethics Committee of Shanghai First Maternity and Infant Hospital of Tongji University (No. 2019-034) and was carried out following the Declaration of Helsinki. Patients provided informed consent before sample collection.

**Table 1.** Patient demographic and obstetrical information.

| Group | m1 |  |  | m2 |  |  |
| --- | --- | --- | --- | --- | --- | --- |
| Patient | PE1 | PE2 | PE3 | Ctrl1 | Ctrl2 | Ctrl3 |
| Maternal age (years) | 33 | 30 | 31 | 40 | 35 | 30 |
| Pre-pregnancy BMI (kg/m<sup>2</sup>) | 20.2 | 17.1 | 21.9 | 22.5 | 21.1 | 21.1 |
| Gravidity | 2 | 1 | 1 | 3 | 2 | 1 |
| Parity | 0 | 0 | 0 | 1 | 0 | 0 |
| Gestational age ( weeks ) | 31+4 | 29+2 | 36+2 | 35 | 34+2 | 35 |
| DM current pregnancy | N | N | N | N | N | N |
| Indications for CS | sPE ,<br>sFGR ,<br>Fetal<br>distress ? | Placental<br>abruption ,<br>sPE | sPE | Vasa<br>previa<br>with<br>placenta<br>previa | Vasa<br>previa,ART | Vasa<br>previa,Marginal<br>placenta previa |
| Birthweight (grams) | 1075 | 1190 | 2905 | 3030 | 2455 | 2430 |
| SGA | Y | Y | N | N | N | N |
| Apgar score ( 1min-5min ) | 8-9 | 7-8 | 9-10 | 8-9 | 9-10 | 9-10 |
| Sex of the baby | M | F | M | M | M | F |

BMI, body mass index; PE, preeclampsia; DM, diabetes mellitus; CS, cesarean section; N, no; Y, yes; SGA, small for gestational age; min, minutes; sPE, severe preeclampsia; sFGR, severe fetal growth restriction; M, male; F, female; ART, Assisted Reproductive Technology

Our processing of self-sequenced single-cell and single-nuclei RNA-sequencing is as described by previous researchers^[31,32]^.The raw data of single-cell self-sequencing were imported using the Seurat package (version 4.3.0) ^[33]^ in R programming language. First, low-quality cells and genes were filtered using the following criteria: 1) removal of genes not detected; 2) retention of cells with the number of expressed genes fluctuating between 200 and 5000; 3) retention of cells with UMI reads less than 20000; 4) retention of cells with a mitochondrial gene percentage below 20%. The "normalizedata" function in the Seurat R (version 4.3.0) package was used for data normalization. After standardization, highly variable genes were identified by balancing the relationship between average expression and dispersion. PCA was then performed, and significant principal components (PCs) were used as input for graph-based clustering. The harmony method was employed to eliminate batch effects from different samples. For clustering, we used the FindClusters function, which is based on a shared nearest neighbor (SNN) modularity-optimized clustering algorithm, producing 16 clusters on 12 PC components at a resolution of 0.6. The "RunUMAP" function was then used for uniform manifold approximation and projection (UMAP). UMAP-1 and UMAP-2 demonstrated cell aggregation. To identify differentially expressed genes between cell clusters, we applied the FindAllMarkers function to the normalized gene expression data, using the default parameters set by Seurat. Subsequently, cell clusters were identified using cell type-specific biomarkers, and the proportions of cell types were calculated and assessed.

### Download and Organization of Routine Transcriptome Data

This part of data used in this study are publicly available and primarily sourced from the GEO (Gene Expression Omnibus, https://www.ncbi.nlm.nih.gov/geo/) database. Expression profile data and clinical data from the GEO database were downloaded using the R package "GEOquery" (version 2.62.2). GSE35574 was sequenced based on the GPL6102 Illumina human-6 v2.0 expression beadchip platform, including 35 samples of Intrauterine Growth Restriction (IUGR, including 8 batch effect technical replicates), 19 samples from preeclampsia, and 40 control placental samples, of which 19 samples were from preeclampsia and 40 from control placentas. GSE24129 was sequenced based on the GPL6244 [HuGene-1_0-st] Affymetrix Human Gene 1.0 ST Array [transcript (gene) version] platform, including 8 samples from preeclampsia, 8 from normotensive pregnancies with Fetal Growth Restriction (FGR), and 8 from pregnancies without FGR, of which 8 samples from preeclampsia and 8 from normotensive pregnancies were included in this study.

In the analysis, we merged the above data to obtain a dataset containing 27 preeclampsia samples and 48 control samples for all analyses. The ComBat method from the R package "sva" (version 3.42.0) ^[34]^ was used to correct for batch effects due to non-biotechnological biases. This facilitated data merging, and principal component analysis (PCA) was employed to check the degree of correction. This study respects the data access policies of each database and primarily relies on public datasets; thus, no additional ethical review is required.

Processing of Single-Cell Sequencing Data

### Deconvolution Analysis

Single-sample gene set enrichment analysis (ssGSEA) (version 2.25.0) is an extension of gene set enrichment analysis (GSEA) that can calculate individual enrichment scores for each pair of samples and gene sets. Each ssGSEA enrichment score represents the degree to which genes in a specific gene set are coordinately upregulated or downregulated in the sample. Based on the list of the top 200 differentially upregulated genes between single-cell types, we calculated the enrichment scores for various cell types in the routine transcriptome data using the ssGSEA algorithm.

### Screening Key Immune Cells

SVM-RFE is a machine learning technique that trains subsets of features from different categories to narrow down the feature set and find the most predictive features. To compute and select linear models while retaining valuable variables, we used the "glmnet" (version 4.1.4) package in R for LASSO regression. Binary distribution variables were then used for LASSO classification, selecting the minimum error to establish the model. Random forest analysis was performed using the "RandomForest" function. The minimum error was selected as the mtry node value, and stable image values were chosen as ntree. We calculated the average decrease in accuracy (MDA) and average decrease in Gini index (MDG) based on feature weights. Then, combining SVM, LASSO regression, and random forest, the most significant feature cells were selected as key immune cells in this study.

### IREA

Immune Response Enrichment Analysis (IREA) is a method proposed by Ang Cui et al. to infer cytokine activity and immune cell polarization in any immune process based on collected gene expression data^[23]^. This method provides a cell type-centered view to determine the polarized states of immune cell types driven by over 66 cytokines, including previously uncharacterized states.

We calculated the cytokines primarily produced in response by key immune cells using IREA based on the upregulated genes in the differential genes (|log2Fold Change|>0.25, p value < 0.05) between m1 (normal preterm birth) and m2 (severe preeclampsia), as well as the polarization state of key immune cells in preeclampsia.

### Cell Communication Analysis and Receptor-Ligand Expression

The CellChat object was created by the "CellChat" R package based on the UMI count matrix for each group (preeclampsia vs. healthy controls) (https://www.github.com/sqjin/CellChat) ^[35]^. Using the "CellChatDB.human" ligand-receptor interaction database as a reference, cell communication analysis was performed using default parameters. The "mergeCellChat" function was used to combine each group’s CellChat objects to obtain a comparison of the total number of interactions and interaction strength. The "netVisual_diffInteraction" function visualized the differences in the number or strength of interactions between different cell types across groups. Finally, we visualized the distribution of signaling gene expression between groups using the "netVisual_bubble" and "netVisual_aggregate" functions.

### Differential Analysis

To clarify the core targets determining the progression of preeclampsia, this study used the R package "limma" (version 3.50.3) ^[36]^ to identify differentially expressed genes (DEGs) between the control group and preeclampsia samples in the merged dataset (27 preeclampsia samples and 48 control samples). The screening criteria for differential genes were P adjust < 0.05, and the resulting differential genes were intersected with the differential genes of key immune cells between preeclampsia and healthy control groups (logfc.threshold > 0.25, p value < 0.05). The intersected genes were considered to play an important role in the occurrence and development of preeclampsia.

### GO/KEGG Pathway Enrichment Analysis

Gene Ontology (GO) ^[37]^ enrichment analysis includes biological processes (Biological Process, BP), molecular functions (Molecular Function, MF), and cellular components (Cellular Component, CC) analysis. The Kyoto Encyclopedia of Genes and Genomes (KEGG)^[38]^is a bioinformatics resource used to mine pathways that are enriched and significantly altered in gene lists. The R package "clusterProfiler (version 4.2.2)" ^[39]^ was applied for GO/KEGG enrichment analysis of the genes of interest (p value < 0.05).

### Machine Learning

To compute and select linear models while retaining valuable variables, we used the "glmnet" (version 2.25.0) package in R for LASSO regression. Binary distribution variables were then used for LASSO classification, selecting the minimum error to establish the model. Random forest analysis was performed using the "RandomForest" function. The minimum error was selected as the mtry node value, and stable image values were chosen as ntree. Based on the average decrease in accuracy (MDA) and average decrease in Gini index (MDG) of feature weights, the top 5 differentially expressed genes were selected as important genes identified by random forest. Then, combining LASSO regression and random forest, we selected feature genes supported by all three machine learning algorithms as key genes in this study.

### GeneMANIA

The GeneMANIA website (http://genemania.org) can predict relationships between functionally similar genes and hub genes, including protein-protein interactions, protein-DNA interactions, pathways, physiological and biochemical responses, co-expression, and co-localization^[40]^. We constructed a protein-protein interaction (PPI) network of key genes through the GeneMANIA website.

### ceRNA Network Construction

Due to the unclear mechanism of competing endogenous RNA (ceRNA) in preeclampsia, we used the miRTarBase (https://mirtarbase.cuhk.edu.cn/~miRTarBase/miRTarBase_2022/php/index.php) ^[41]^, starbase2.0 (https://starbase.sysu.edu.cn/starbase2/index.php)^[42]^, and miRDB (https://mirdb.org/index.html) databases to perform reverse prediction of microRNAs for key genes to predict lncRNAs that co-target key genes and ultimately construct a ceRNA network.

### RBP-mRNA Network Construction

This study used the widely used open-source platform StarBase (https://starbase.sysu.edu.cn/tutorialAPI.php#RBPTarget) to analyze lncRNA interactions, using CLIP-seq, degradome-seq, and RNA-RNA interaction data to study the association between mRNA and RBP (RNA-binding proteins) expression. In preeclampsia, p value < 0.05, clusterNum ≥ 30, and clipExpNum ≥ 5 were defined as cutoff criteria for identifying key mRNA-RBP pairs. Subsequently, the RBP-mRNA network was constructed using Cytoscape.

### Drug-Gene Interaction Analysis

The Drug-Gene Interaction Database (DGIdb, https://dgidb.org) is a publicly accessible resource that compiles records of genes or gene products, drugs, and drug-gene interactions to facilitate hypothesis generation and discovery for clinicians and researchers ^[43]^. We utilized the DGIdb database to predict potential drugs or small molecular compounds that interact with key genes.

### Statistical Analysis

This study used R software v4.1.2 for statistical analysis. Spearman correlation tests were used to infer the correlation between two parameters. Wilcoxon tests were used to compare differences between two groups. A p-value less than 0.05 was considered statistically significant.

## Results

The flowchart of this study is shown in Figure 1.

### Single-Cell Dimensionality Reduction Clustering and Annotation

We demonstrated the complex cellular environment in preeclampsia through dimensionality reduction clustering annotation of single-cell sequencing data. After preliminary quality control assessment and doublet removal, we successfully filtered 63,387 cells from single cells. Next, we used clustering methods to divide all cells into 16 clusters (Figure 2A) and annotated these cell types based on the gene expression characteristics of each cluster, combined with cell-specific biomarkers (Table S1). As shown in Figure 2B, we identified 9 major cell types in preeclampsia, including Syncytiotrophoblast, SMC, Fibroblast, Endothelial, Villous cytotrophoblast, Macrophage, Lymphocyte, Neuron, and Extravillous trophoblast. Through dot plots, we visually displayed the annotated genes for each cell type (Figure 2C). Meanwhile, in Figure 2D, we presented the proportion distribution of different cell types in m1 and m2 samples, where Syncytiotrophoblast and Lymphocyte showed changes in proportion before and after the occurrence of mild to severe preeclampsia, indicating that these cells play an important role in the progression of preeclampsia.

**Figure 2.**
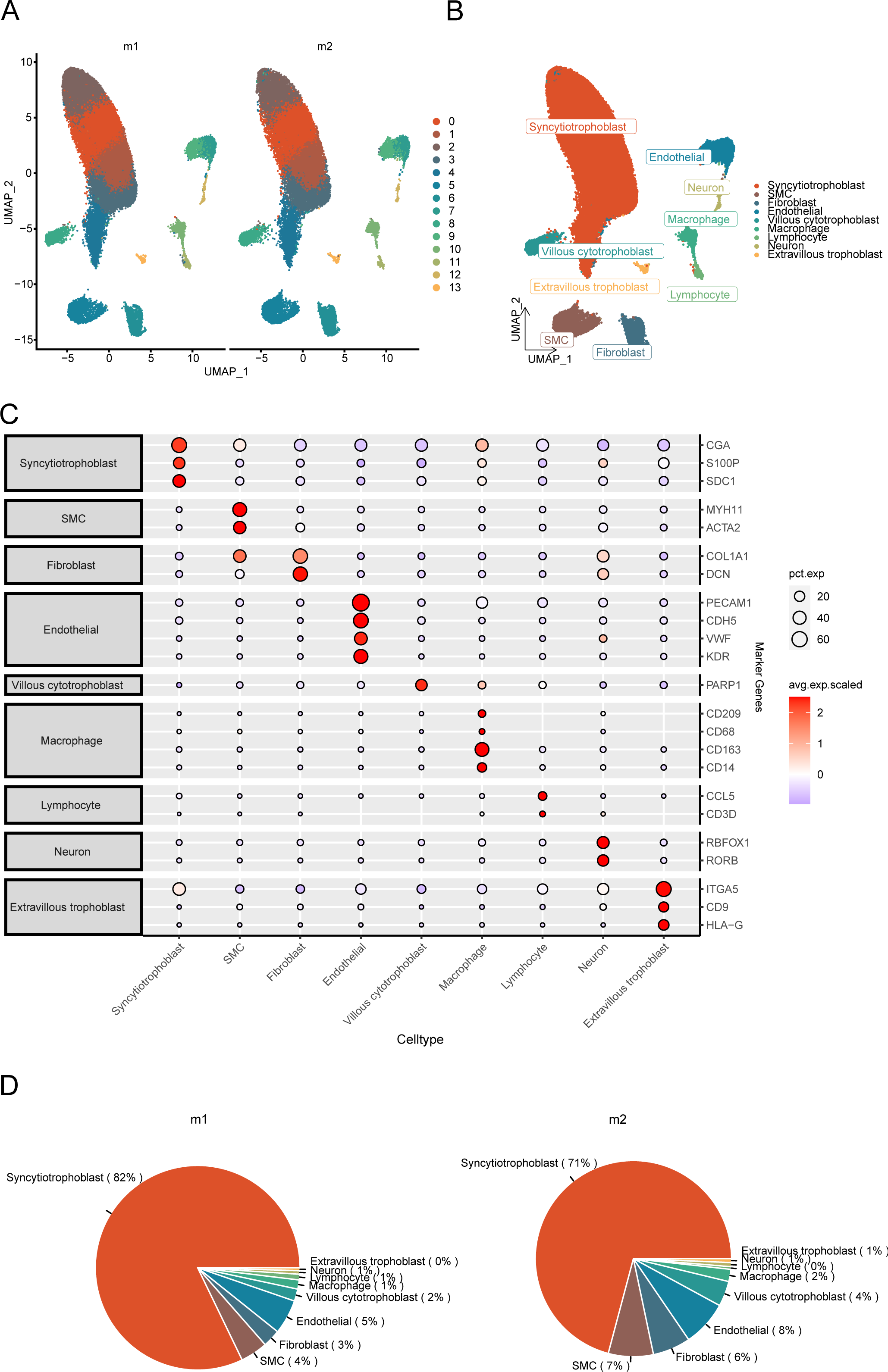
Annotation and visualization of the cellular environment in preeclampsia. (A) UMAP plot showing the distribution of 16 cell subpopulations in the m1 and m2 groups. (B) UMAP plot displaying the annotation results and distribution of 9 major cell types in preeclampsia. (C) Expression of marker genes in each cell type. (D) Pie chart showing the proportion of various cell types in the normal preterm (m1) group and the severe preeclampsia (m2) group. Left: m1 group. Right: m2 group.

### Lymphocyte Re-Clustering

We used clustering methods to re-cluster lymphocytes into 7 clusters (Figure S1A) and annotated these cell types based on the gene expression characteristics of each cluster, combined with cell-specific biomarkers (Table S2). As shown in supplementary Figure S1B, we identified 7 cell subtypes in preeclampsia, including CD8+ T cells, NK cells, Th cells, CD4+ T cells, gamma delta T, B cells, and Treg. Through dot plots, we visually displayed the annotated genes for each cell type (Figure S1C). Meanwhile, in supplementary Figure S1D, we presented the proportion distribution of different cell subtypes in the normal preterm (m1) group and the severe preeclampsia (m2) group, where NK cells and CD4+ T cells showed changes in proportion before and after the occurrence of mild to severe preeclampsia, indicating that these cells play an important role in the occurrence of preeclampsia.

### Deconvolution of Single-Cell Data and Correlation Analysis of Immune Cells

Combining the re-clustered cell subtypes with the aforementioned 9 cell types, we obtained a total of 15 cell types. Using the top 200 most significant genes from the differentially expressed genes between cell types (|log2FC|>0.25, p.value <0.05, Table S3) for deconvolution. By plotting box plots, we demonstrated that the infiltration levels of cell types in single-cell data also showed significant differences before and after the occurrence of PE (Figure 3A). A bubble plot was used to show the correlation between cells, revealing that most cell types exhibited positive correlations (Figure 3B), indicating that these immune cells play important roles in the occurrence and further progression of PE.

**Figure 3.**
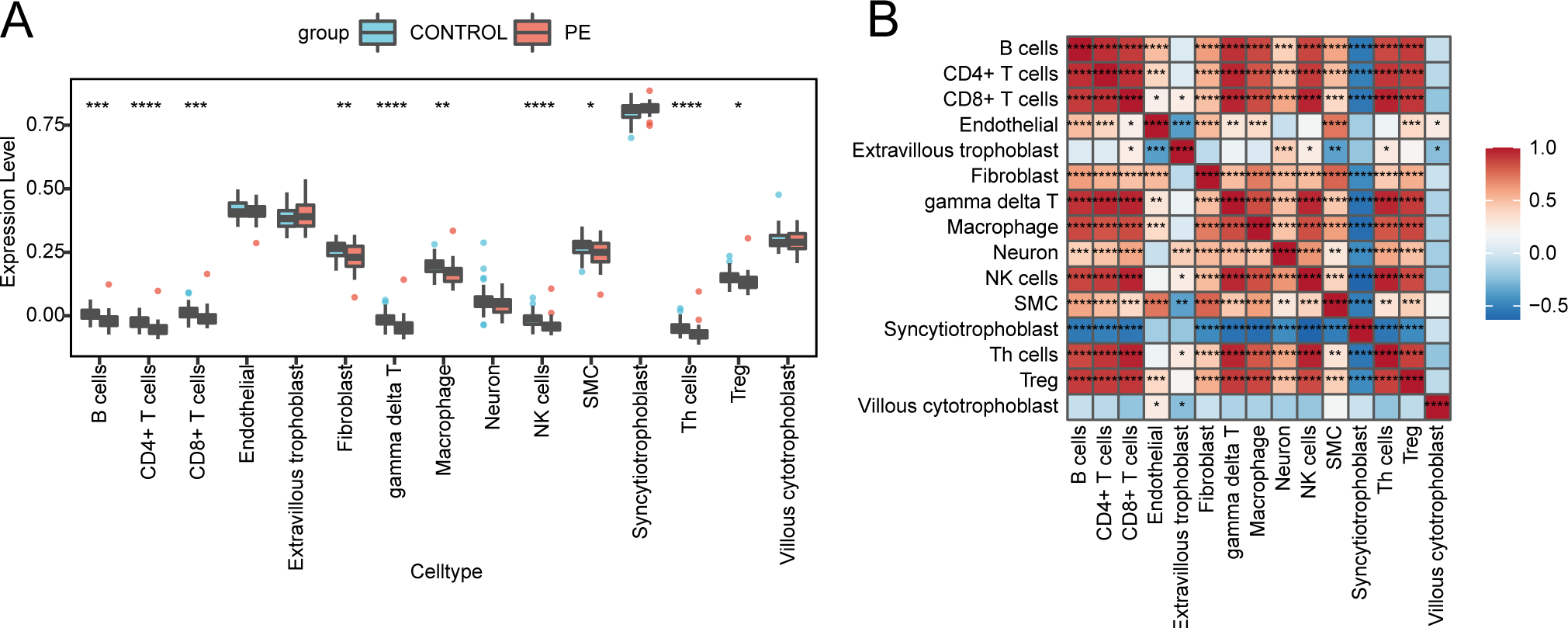
Infiltration levels of cell types between m1 and m2 groups in scRNA-seq data. (A) Box plot showing the estimated proportions of cell type infiltration between the PE and control groups after deconvolution. (B) Heatmap displaying the correlation between cells. Asterisks indicate P values: ****p < 0.0001, ***p < 0.001, **p < 0.01, *p < 0.05.

### Screening Key Immune Cells Using Machine Learning Algorithms

We further utilized LASSO regression and random forest algorithms to screen for key cells. Through LASSO regression analysis, we identified 2 key cells (Figure 4A&4B). Using the random forest algorithm, we selected the intersection of the top 5 immune cells from both methods based on feature weights MDA and MDG as key cells (Figure 4C&4D). Next, we used SVM-RFE to screen for 2 types of cells (Figure 4E&4F). The intersection of key cells detected by the three methods yielded the most critical immune cell, CD4+ T cells (Figure 4G). We analyzed the diagnostic performance of multiple immune cells using ROC curves, finding that the most critical immune cell, CD4+ T cells, had the highest AUC value of 0.7878, indicating good diagnostic efficacy for PE (Figure 4H).

**Figure 4.**
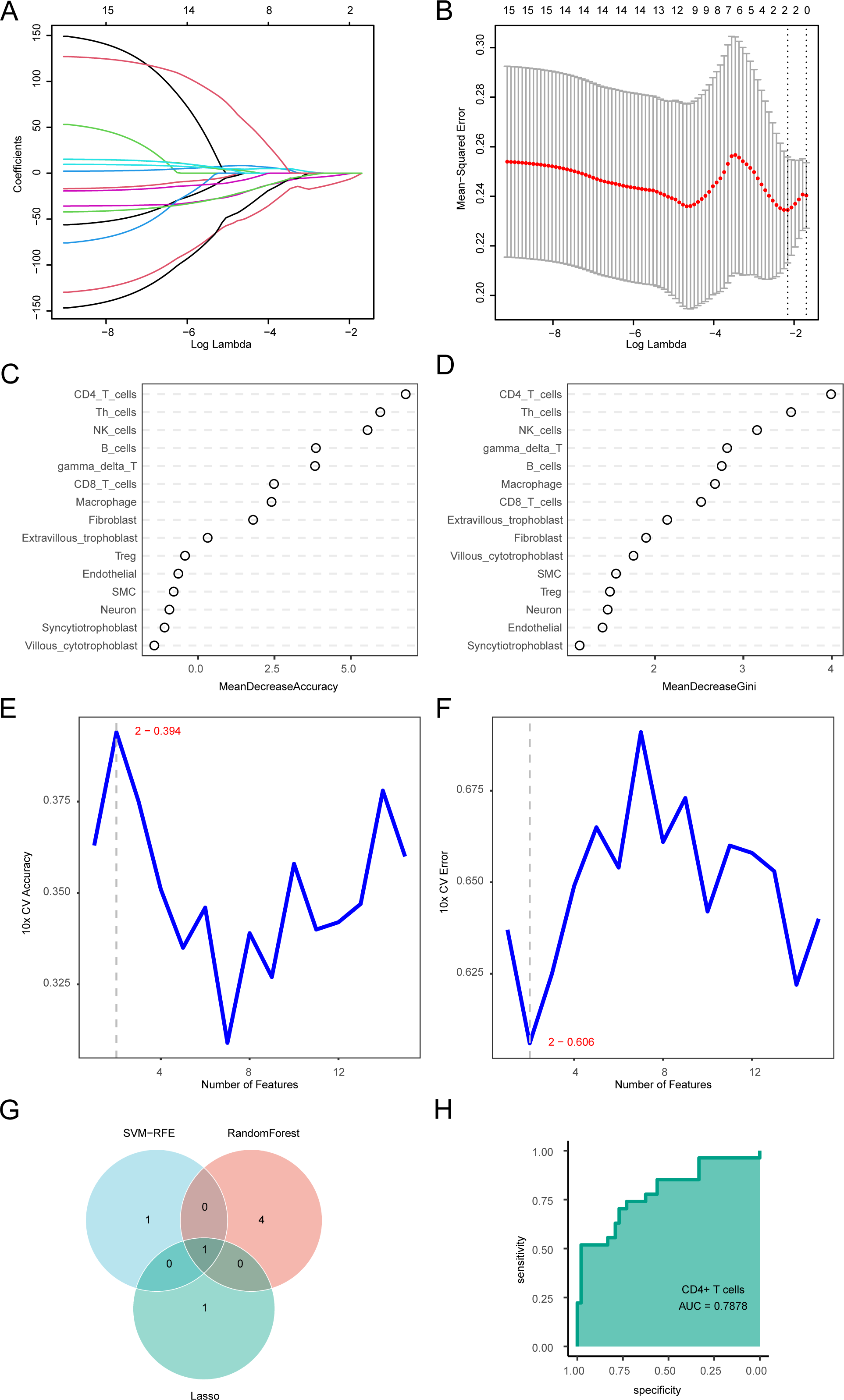
Selection of key immune cells in the process of preeclampsia using machine learning methods. (A) LASSO regression trajectory of independent variables, with the x-axis representing the logarithm of the independent variable lambda and the y-axis representing the coefficients that can be independently obtained. (B) Confirmation intervals for each lambda in LASSO regression. (C) Accuracy importance measure of potential targets. (D) Gini importance measure of potential targets. (E) SVM accuracy curve. (F) SVM error rate curve. (G) Venn diagram showing the intersection of three machine learning algorithms. (H) ROC curve of immune cells CD 4+ T cells.

### Immune Response Enrichment Analysis of Key Immune Cells

To explore the immune status of key immune cells in preeclampsia, we conducted differential analysis between the m1 group and m2 group for key immune cells, identifying a total of 143 DEGs, with statistically significant differences between the two groups (|log2FC| >0.25, p_value < 0.05, Table S4). The heatmap detailed the expression of the 10 most significantly different feature genes in key immune cells (Figure 5A). Subsequently, we performed IREA analysis based on the significantly upregulated genes in severe preeclampsia. For the key immune cell CD4+ T cells, the cell polarization radar plot (Figure 5B, Table S5) indicated that CD4+ T cells were primarily in the T4-c polarized state in the severe preeclampsia group; the cytokine enrichment plot showed that CD4+ T cells mainly enriched cytokines such as IL-1a and IL-1b in the severe preeclampsia group (Figure 5C, Table S6). IL-1α and IL-1β may exacerbate the pathological process of preeclampsia by promoting inflammatory responses. They can activate immune cells and increase vascular inflammation, consistent with the endothelial dysfunction and vascular inflammation observed in preeclampsia. Additionally, these cytokines may directly participate in pathological changes in the placenta by affecting placental function and the proliferation and invasion of trophoblasts. Furthermore, the increased expression of PTX3 induced by IL-1β may play a role in the development of preeclampsia. PTX3, as a pro-inflammatory marker, may inhibit trophoblast function through its excessive expression in the trophoblast layer, thereby affecting normal placental development and function ^[44]^.

**Figure 5.**
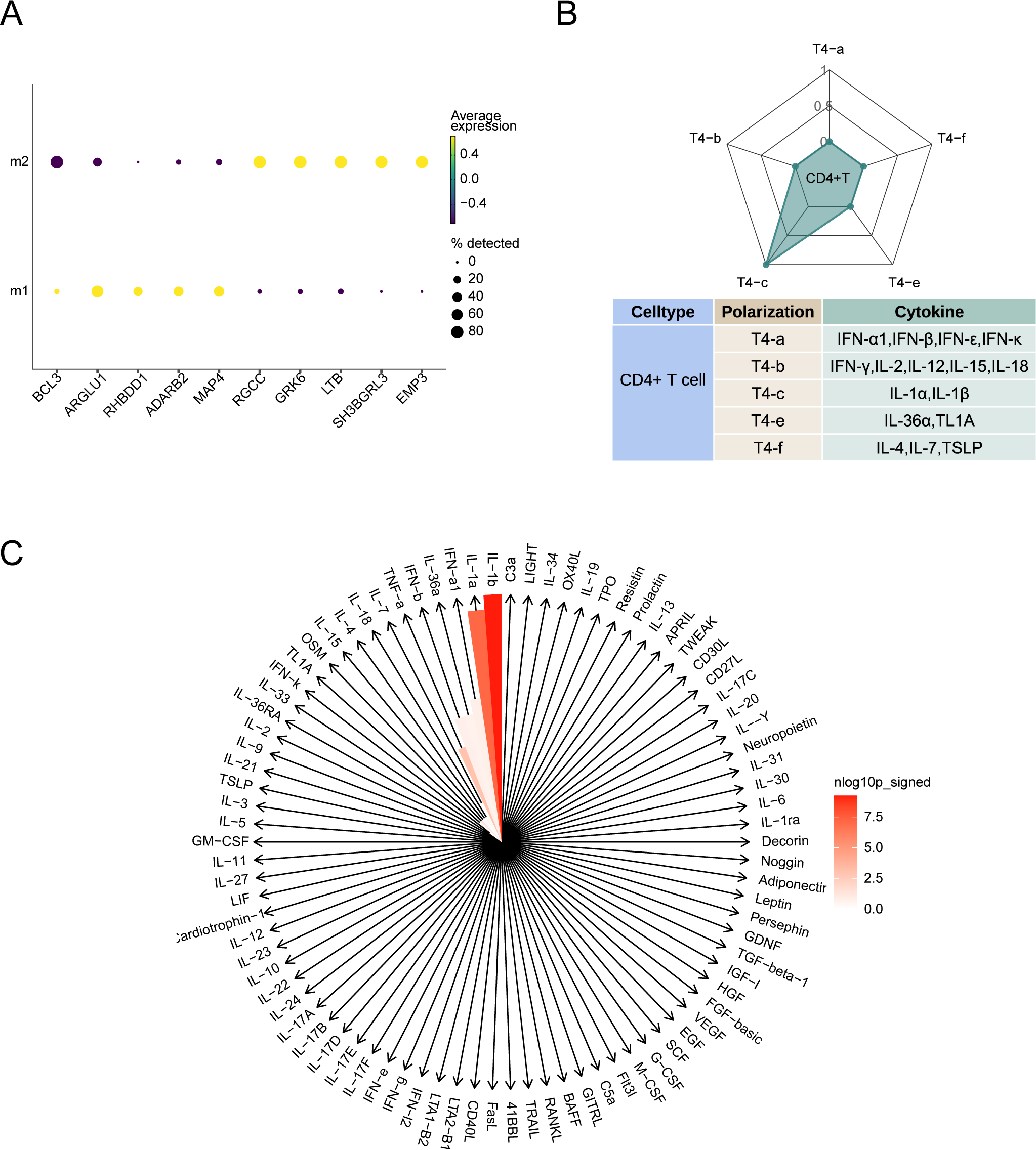
Analysis of immune response in CD4+ T cells. (A) Dot plot showing 10 significantly differentially expressed genes in key immune cells between the m1 and m2 groups. (B) Radar plot of cell polarization of CD4+ T cells in severe preeclampsia. (C) Cytokine enrichment plot of CD4+ T cells in severe preeclampsia.

### Cell Communication

Previous analyses have identified the important role of CD4+ T cells in the occurrence of preeclampsia. We further explored the significant changes in cell communication involving CD4+ T cells as participants to explain the main reasons for the functional realization of CD4+ T cells. First, we used the R package "Cellchat" to reveal the changes in cell crosstalk between the normal preterm and severe preeclampsia groups. Compared to normal preterm, both the total number and strength of interactions decreased (Figure 6A), with most interactions between all cells showing reduced quantity and weakened intensity (Figure 6B). We then further compared the overall signaling patterns between tissues from normal preterm and severe preeclampsia. The overall signaling patterns of the normal preterm and severe preeclampsia groups are clearly displayed in Figure 6C. For example, the signal intensities of EGF, OSM, and SPP1 originating from CD4+ T cells changed in severe preeclampsia, indicating that this pathway may be related to the changes in the PE process.

**Figure 6.**
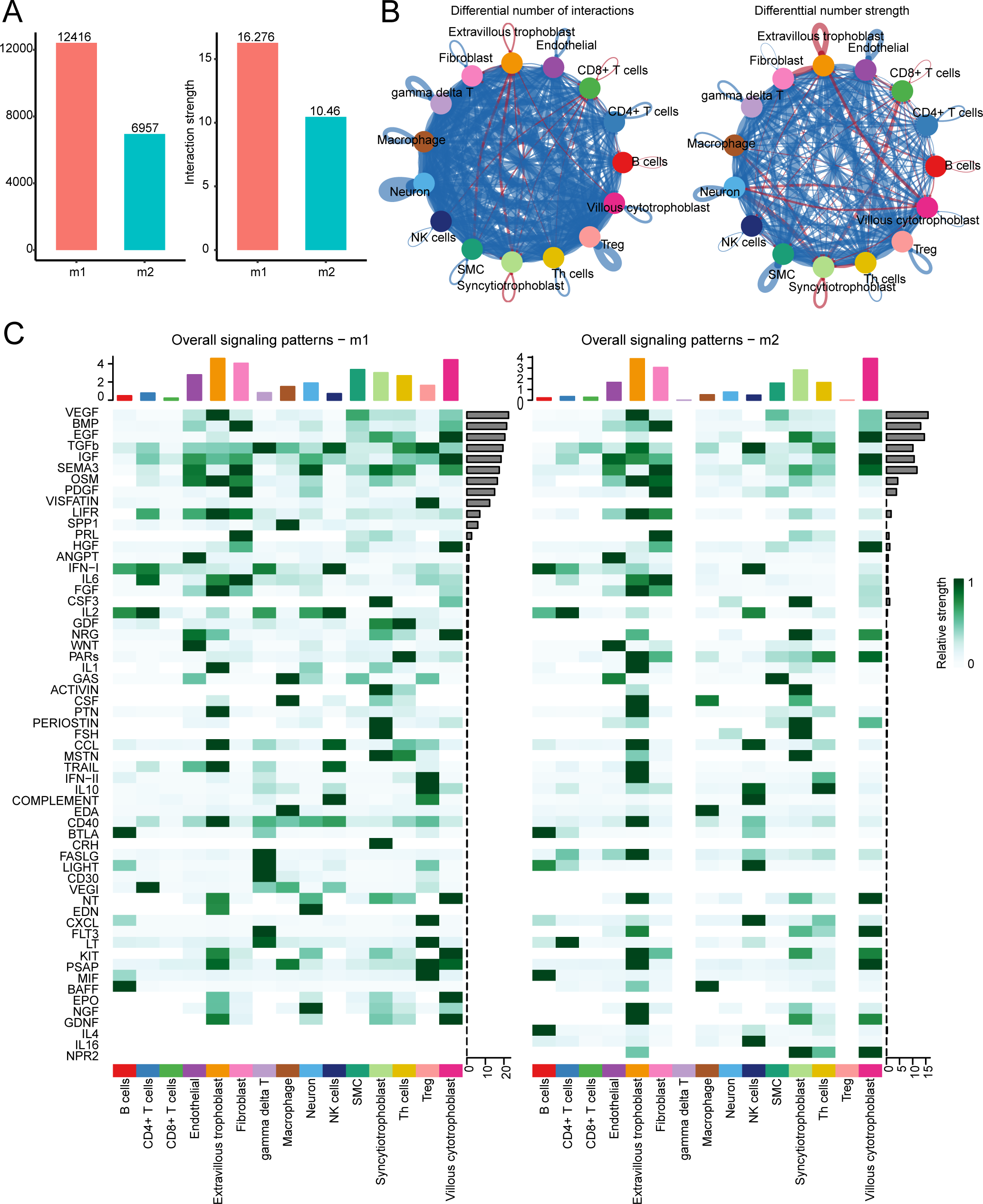
Number and intensity of cell communications between m1 and m2 groups. (A) Bar chart showing the total number and intensity of interactions between all cell types in single cells of the m1 and m2 groups. (B) Network diagram showing intergroup changes in the number and intensity of interactions between various cell types in the m1 and m2 groups, with red lines indicating an increase in total communication quantity and intensity in severe preeclampsia compared to the control group, and blue indicating a decrease, left: communication quantity, right: communication intensity. (C) Heatmap depicting the signaling pathways contributing the most to the overall signaling pathways in the m1 and m2 groups.

Additionally, we analyzed receptor-ligand pairs that may regulate communication between key immune cells and other cells. The SPP1 signaling pathway is an important signaling pathway in preeclampsia. The SPP1 ligand from Macrophage binds to the corresponding receptor on key immune cells CD4+ T cells, and this interaction was observed and significantly increased in the normal preterm group, but not found in the severe preeclampsia group, indicating that SPP1 is a pathway with reduced signal strength in severe preeclampsia (Figure 7A). The violin plot shows the levels of SPP1 signaling pathway receptors and ligands expressed by different cell types in the normal preterm and severe preeclampsia groups (Figure 7B). In contrast, there is a significant trend of reduced expression of SPP1 receptors in CD4+ T cells in the severe preeclampsia group, explaining the main reason for the decreased SPP1 communication between macrophages and CD4 T cells.

**Figure 7.**
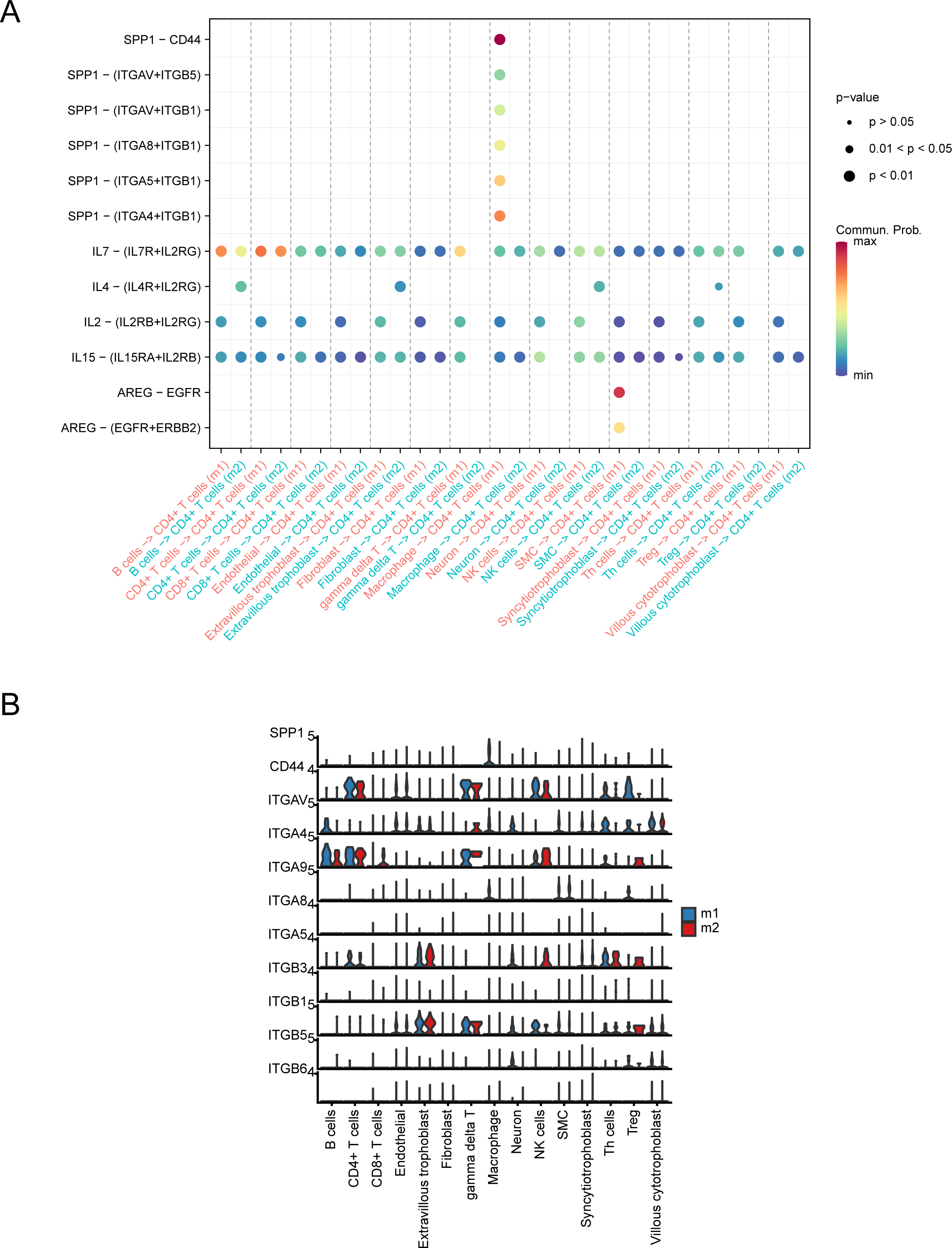
Signaling communications that significantly changed between m1 and m2 groups, major ligand-receptor pairs, and their expression. (A) In the severe preeclampsia group, the ligand-receptor pairs with increased communication intensity between CD4+ T cells and other cell types. (B) Violin plot showing the expression distribution of SPP1 signaling ligands in the m1 and m2 groups.

### Obtaining Intersection Genes and Their Enrichment Analysis

To further explore the key targets regulated by key immune cells that participate in the occurrence and development of preeclampsia, we integrated single-cell and transcriptome data to find the decisive gene group. By comparing the preeclampsia group (PE) and control group from the transcriptome analysis, a total of 1201 differentially expressed genes (DEGs) were identified, with statistically significant differences between the two groups (FDR adjusted P value <0.05). In PE samples, 532 genes were upregulated, and 669 genes were downregulated (Table S7). The heatmap displays the top 5 upregulated genes (CSNK1E, PPP1R12C, ENG, HTRA4, LMBR1L) and the top 5 downregulated genes (NDUFA12, CTTNBP2NL, GABARAPL2, SNAI2, B3GNT2) ranked by P value (Figure 8A). Based on the important role of CD4+ T cells in preeclampsia, we intersected the significantly different feature genes of key immune cells between the m1 group and m2 group with the transcriptome DEGs, resulting in 9 intersected genes (Figure 8B, Table S8), which are considered to be differentially expressed in CD4+ T cells and play important roles in disease occurrence.

**Figure 8.**
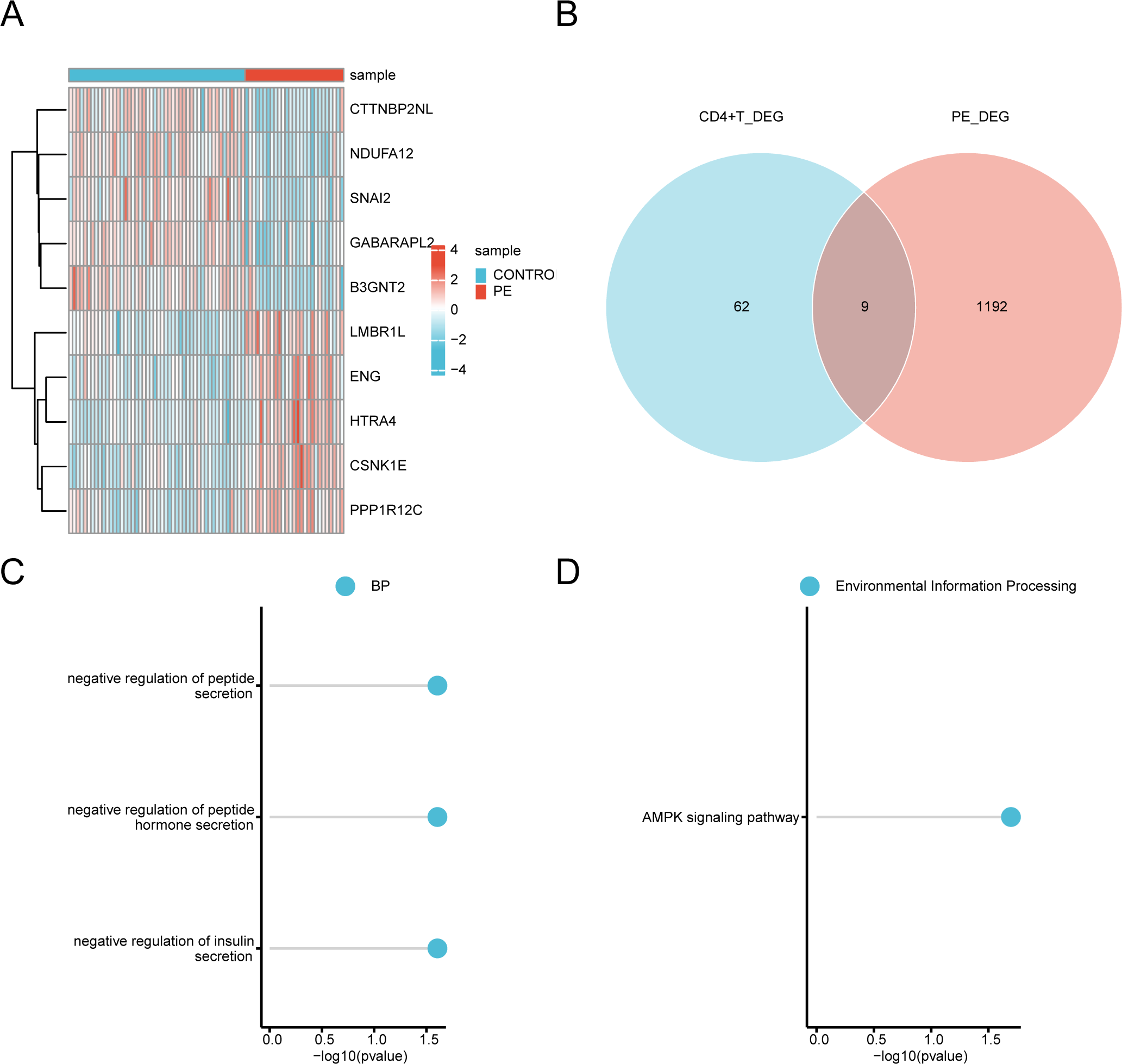
Acquisition and enrichment analysis of major targets in preeclampsia. (A) Heatmap of the top ten differentially expressed genes between preeclampsia and control groups. (B) Venn diagram showing the acquisition of intersecting genes. (C) GO enrichment analysis of intersecting genes. (D) KEGG enrichment analysis of intersecting genes.

To investigate the biological functions related to these important targets, we conducted GO terms (Table S9) and KEGG pathway (Table S10) enrichment analyses. GO results showed that these genes were enriched in negative regulation of insulin secretion, negative regulation of peptide hormone secretion, and negative regulation of peptide secretion (biological process, BP) (Figure 8C). These results suggest that the main targets mediated by CD4 T cells play roles in the development of preeclampsia by affecting placental function, hormone balance, and vascular health.

Enriched KEGG pathways include the AMPK signaling pathway (Environmental Information Processing) (Figure 8D), indicating that these pathways may be potential mechanisms through which these important targets exert their effects in preeclampsia.

### Screening Key Genes Using Multiple Machine Learning Algorithms

Based on the intersected genes, we further utilized LASSO regression and random forest algorithms to screen for the most critical genes among them. Through LASSO regression analysis, we identified 9 key differential genes (Figure 9A&9B). Using the random forest algorithm, we selected the top 5 genes as related differential genes based on feature weights MDA and MDG (Figure 9C&9D). Finally, the intersection of key differential genes detected by the two methods yielded 4 most critical differential genes for subsequent analysis, including LCP1, PFKL, TSPYL2, and SREBF1 (Figure 9E, Table S11).

**Figure 9.**
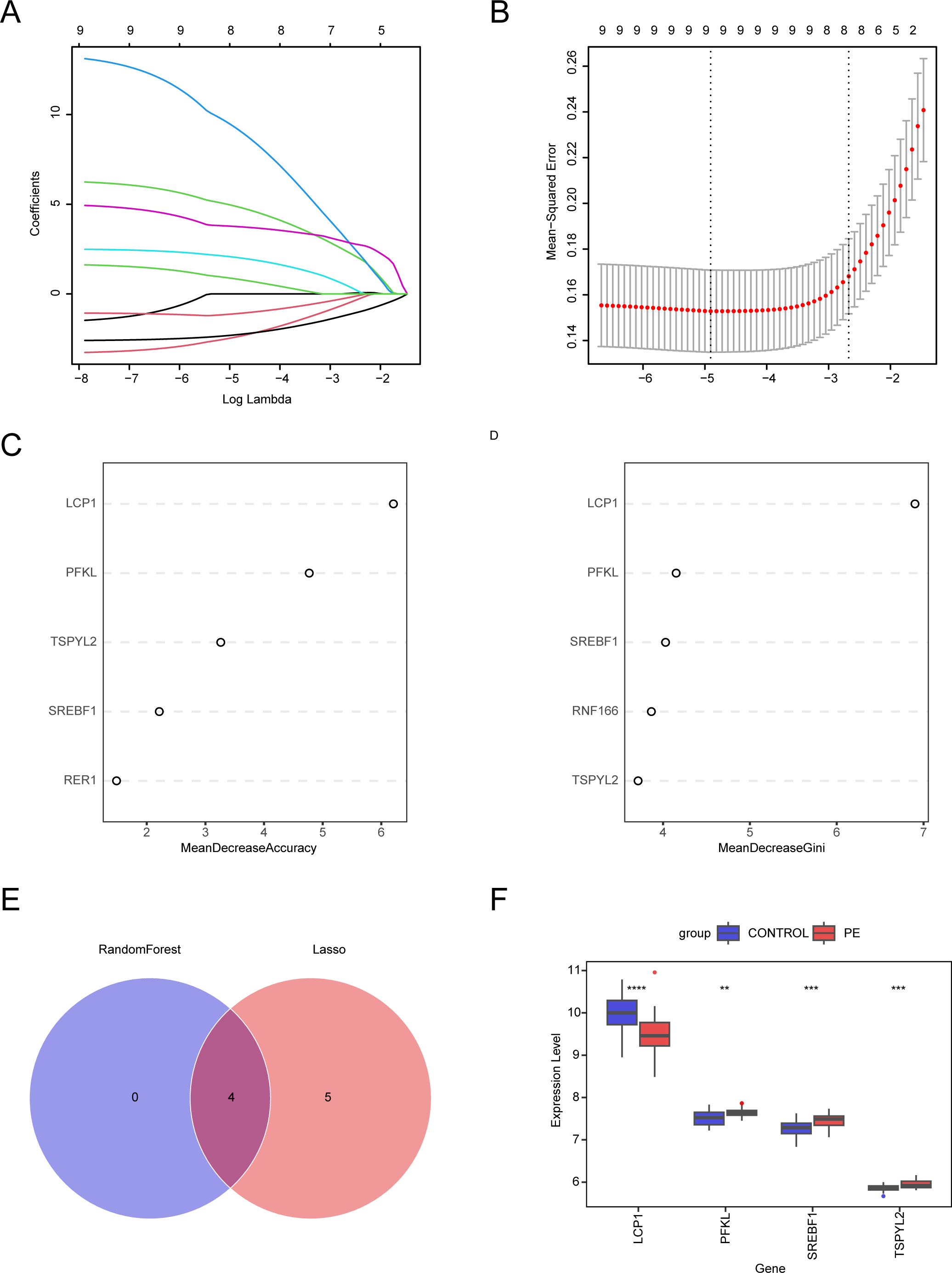
Selection of candidate diagnostic biomarkers for PE progression using machine learning methods. (A) LASSO regression trajectory of independent variables, with the x-axis representing the logarithm of the independent variable lambda and the y-axis representing the coefficients that can be independently obtained. (B) Confirmation intervals for each lambda in LASSO regression. (C) Top 5 DEGs ranked by MDG importance in the random forest algorithm. (D) Top 5 DEGs ranked by MDG importance in the random forest algorithm. (E) Venn diagram showing the intersecting genes from two machine learning algorithms. (F) Box plot of hub gene expression patterns between PE and Control groups.

Additionally, we explored the expression patterns of the 4 hub genes between the PE group and Control, with results shown in Figure 9F, indicating significant expression differences for the 4 hub genes between the two groups.

### Enrichment Analysis of HALLMARK Pathways and Correlation Exploration

To explore the roles of common pathways in the development of diseases in the occurrence of preeclampsia and to investigate the correlation between hub genes and these pathways, we further used ssGSEA to study the differences in 50 HALLMARK signaling pathways between the preeclampsia group and the control group (Table S12). In patients with preeclampsia, significantly downregulated pathways included: HALLMARK_ADIPOGENESIS, HALLMARK_BILE_ACID_METABOLISM, HALLMARK_HEME_METABOLISM, HALLMARK_MYC_TARGETS_V1, HALLMARK_PEROXISOME, HALLMARK_TGF_BETA_SIGNALING, and HALLMARK_UV_RESPONSE_DN (Figure 10A).

**Figure 10.**
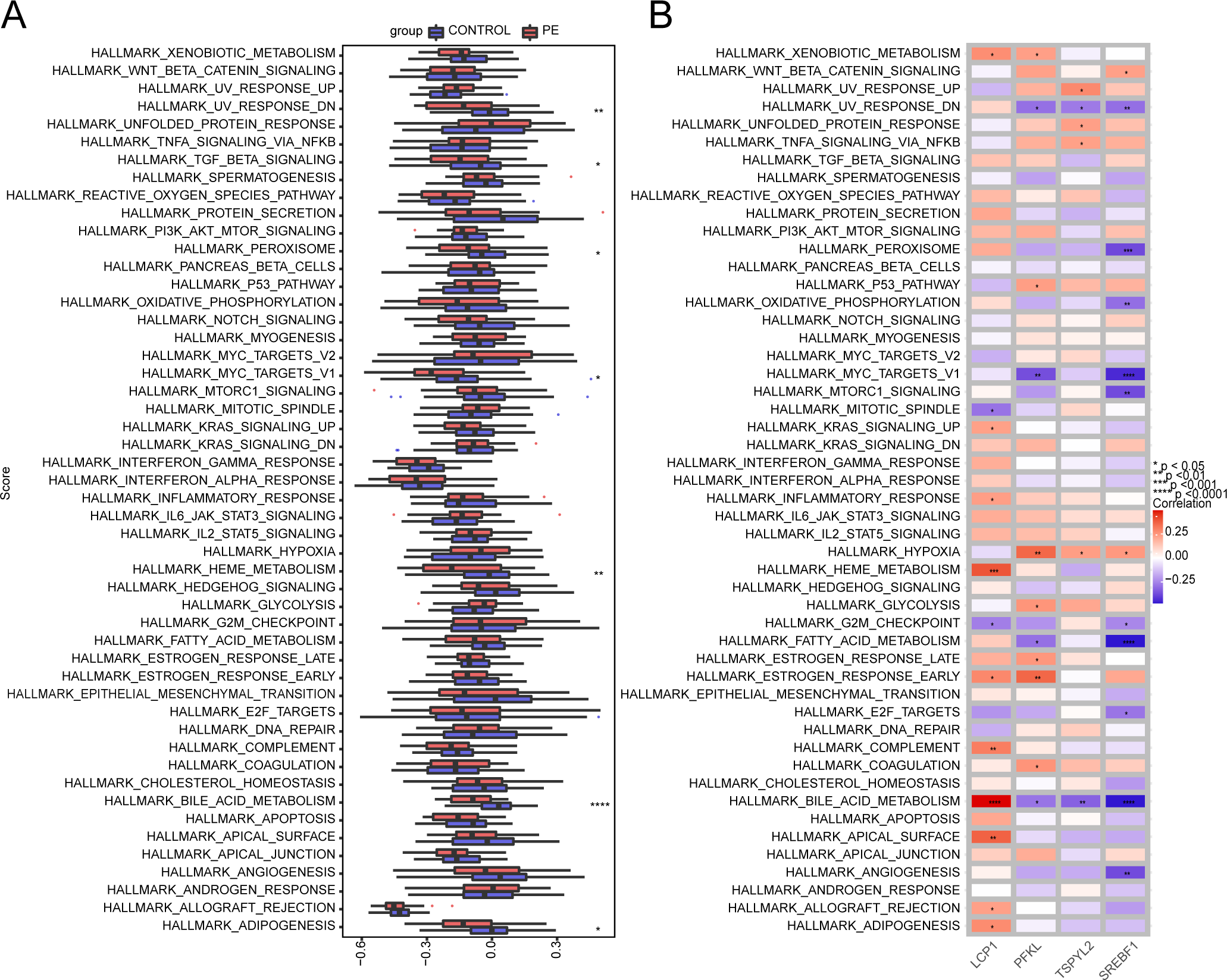
Comparison of 50 HALLMARK signaling pathways between PE and CONTROL groups. (A) Box plot of 50 HALLMARK signaling pathways between PE and CONTROL groups. (B) Heatmap of the correlation between hub genes and 50 HALLMARK signaling pathways. ****p < 0.0001, ***p < 0.001, **p < 0.01, *p < 0.05.

Furthermore, we also explored the correlation analysis between hub genes and signaling pathways. We selected the significantly correlated (p < 0.05) signaling pathways (Figure 10B, Table S13). We found that, except for LCP1, the other 3 hub genes were significantly negatively correlated with pathways such as HALLMARK_BILE_ACID_METABOLISM and HALLMARK_UV_RESPONSE_DN, while showing significant positive correlation with HALLMARK_HYPOXIA. LCP1 was significantly positively correlated with pathways such as HALLMARK_BILE_ACID_METABOLISM, HALLMARK_APICAL_SURFACE, and HALLMARK_HEME_METABOLISM. This suggests that hub genes may participate in the occurrence of preeclampsia by influencing these pathways.

### Interaction Analysis of Hub Genes

We created a PPI network for hub genes using the GeneMANIA database, showing interactions among the 4 hub genes (Figure 11A). To further investigate the functions of feature genes, we conducted GO and KEGG analyses on a total of 24 genes (including 4 hub genes, 20 related genes, and 241 connections, Table S14). GO enrichment results showed that these genes were significantly enriched in biological processes such as "glycerol ether biosynthetic process," "mRNA transcription by RNA polymerase II," and "ether lipid biosynthetic process" (biological process, BP); and in molecular functions such as "negative regulation of leukocyte mediated immunity" (Figure 11C, Table S15). KEGG analysis results indicated that the main enriched pathways were "Glycolysis / Gluconeogenesis," "Carbon metabolism," and "Biosynthesis of amino acids" (Figure 11D, Table S16). We also explored the correlations among the 4 hub genes, finding that LCP1 was significantly negatively correlated with TSPYL2, while SREBF1 was significantly positively correlated with PFKL and TSPYL2 (Figure 11B).

**Figure 11.**
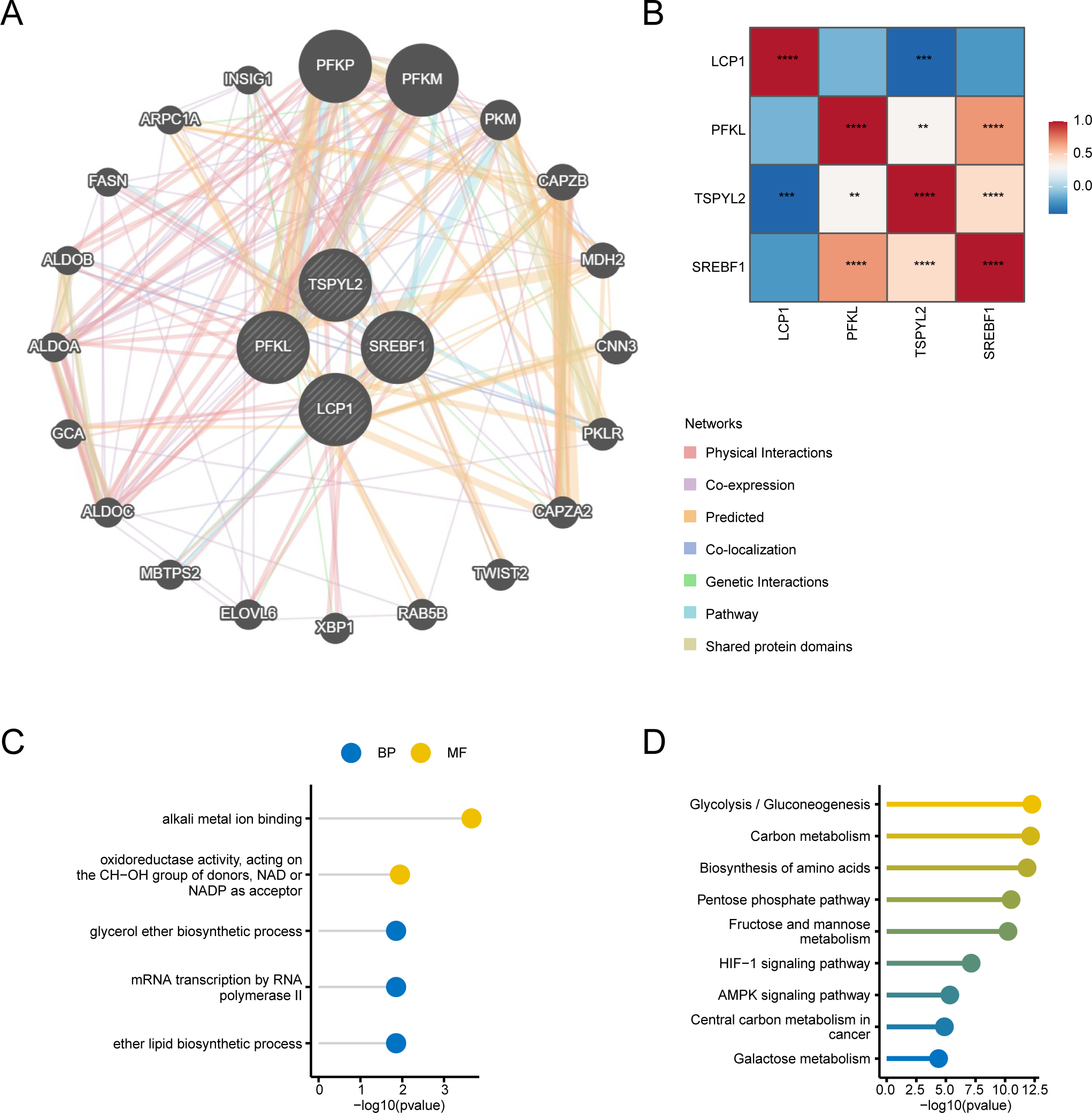
Interaction analysis of hub genes. (A) Gene co-expression network diagram. (B) Correlation between the 4 hub genes. (C) GO enrichment analysis of co-expressed genes. (D) KEGG enrichment analysis of co-expressed genes.

### Drug Prediction of Hub Genes in DGIdb and Construction of ceRNA Network and RBP Network

DGIdb was used to identify potential sensitive drugs or molecular compounds. We performed drug sensitivity analysis on the 4 hub genes in DGIdb and found that only the hub genes SREBF1 and TSPYL2 showed correlations with drugs. As shown in the drug-gene interaction network (Figure 12A, Table 2), a total of 6 drugs or molecular compounds were identified, which have varying degrees of regulatory effects on hub genes. Among the 6 drugs, 4 can be queried for their 2D structures via PubChem (Figure 12B-12E).

**Figure 12.**
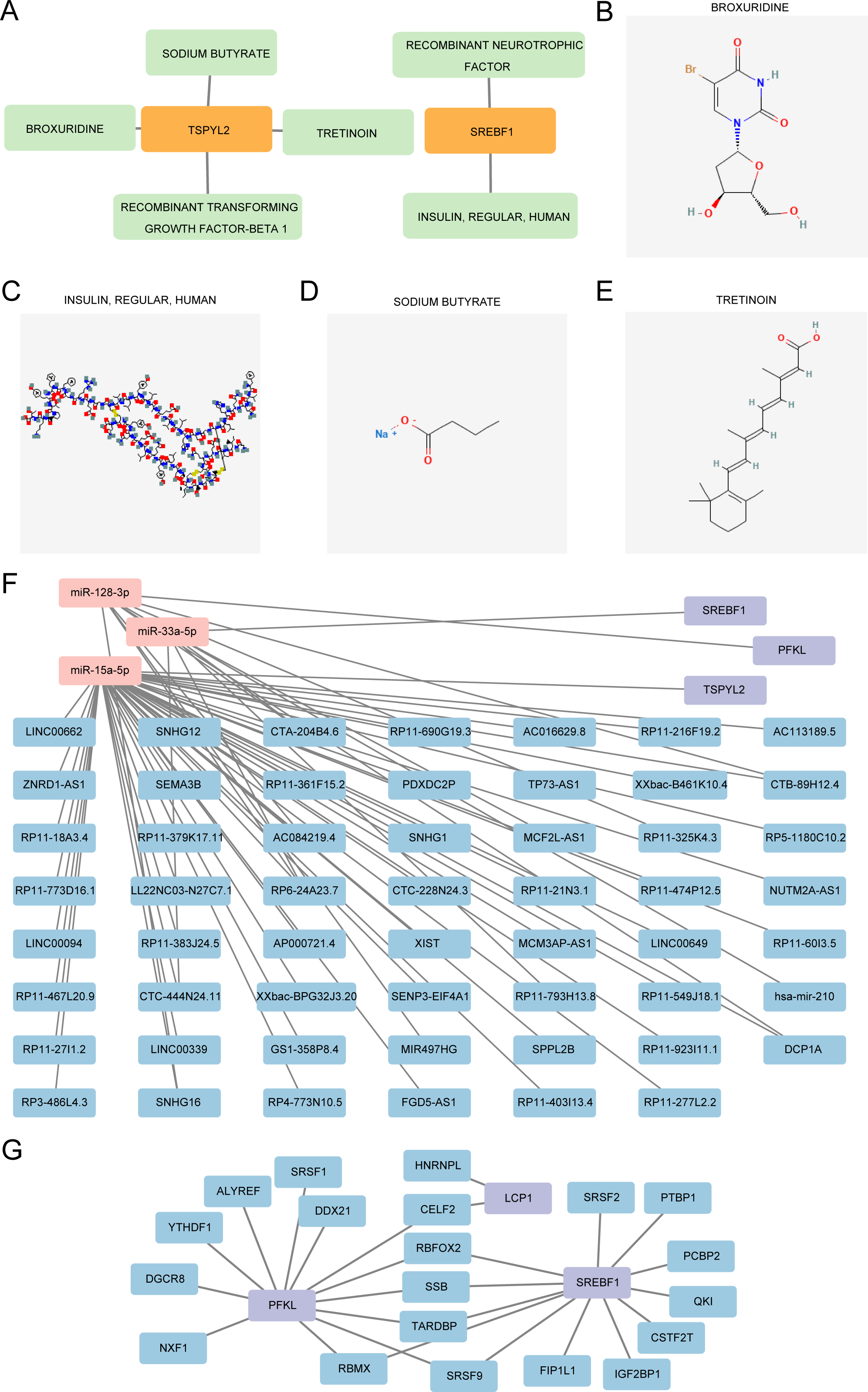
DGIdb drug prediction for hub genes and construction of ceRNA and RBP networks. (A) mRNA-drugs interaction network of hub genes. Green rectangles represent drugs, orange rectangles represent hub genes. 2D structures of BROXURIDINE (B), INSULIN, REGULAR, HUMAN (C), SODIUM BUTYRATE (D), SODIUM BUTYRATE (E). (F) ceRNA network. Purple rectangles represent mRNA, pink rectangles represent miRNA, blue rectangles represent lncRNA. (B) RBP-mRNA regulatory network. Blue rectangles represent RBP, purple rectangles represent mRNA.

**Table 2.**
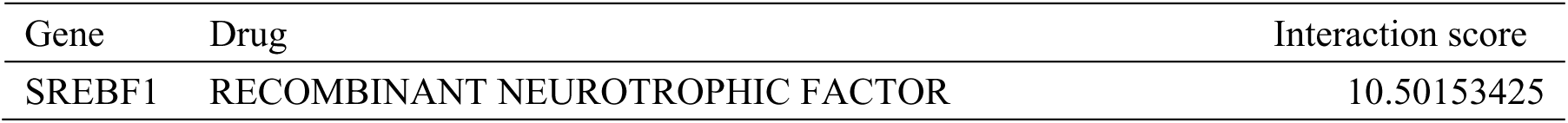
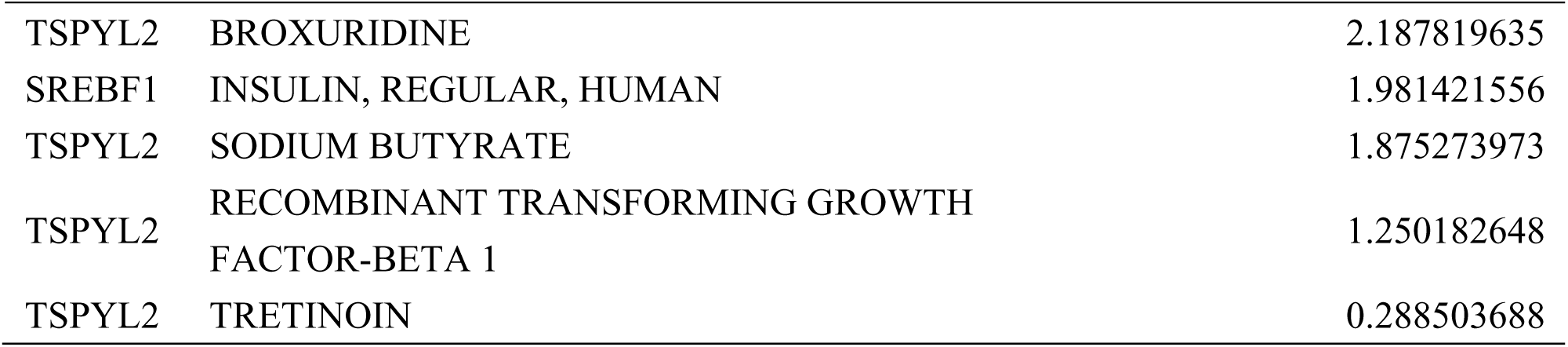
Drug-gene interaction information.

Additionally, to clarify the potential regulatory mechanisms of the 4 hub genes in preeclampsia, we constructed an mRNA–miRNA–lncRNA interaction network, successfully building the mRNA–miRNA–lncRNA interaction network for all hub genes except LCP1 (Figure 12F, Table S17).

Since RBPs (RNA-binding proteins) bind to mRNA, we utilized the StarBase online database to search and download the corresponding mRNA/RBP pairs for the 4 hub mRNAs. Except for TSPYL2, the other hub genes have corresponding RBPs. Based on the relationships provided by the online dataset, we constructed an RBP-mRNA network, which includes 23 nodes, 20 RBPs, 3 mRNAs, and 26 edges (Figure 12G, Table S18).

## Discussion

Although previous researches^[28,45,46]^ indicate the immune system exerts a crucial function in the progression of PE, very little was found in the literature on the question of how exactly immune cells and cytokines work. An initial objective of the study was to identify early detection targets and potential immunological treatments for PE since it remains a significant cause of maternal and fetal morbidity and mortality, with current treatment strategies limited to symptom management and delivery. Using scRNA-seq, we characterized 63,387 placental cells from sPE patients and matched controls to explore immunological alterations in PE. The results of this study indicate that CD4+ T cells exhibit significant alterations in their abundance and functional states during the progression of SPE, particularly their polarization towards a T4-c state characterized by the production of pro-inflammatory cytokines such as IL-1α and IL-1β. Cell–cell communication analysis uncovered alteration in SPP1 signaling between macrophages and CD4+T cells, suggesting immune response dysregulation. Furthermore, machine-learning algorithms identified four hub genes. Six small-molecule drugs were predicted to have therapeutic potential for PE. Our study indicated that the prevention and treatment approach for PE should maybe gradually transition to regulating the immune response and subduing the inflammatory reaction. These nascent therapeutics are anticipated to furnish patients with PE with more efficacious and perdurable resolutions to mitigate symptoms and improve maternal and fetal outcomes.

The findings of this study highlight the significant role of CD4+ T cells in the pathogenesis of PE, particularly their polarization towards a T4-c state characterized by the production of pro-inflammatory cytokines such as IL-1α and IL-1β. In accordance with the present results, previous studies have demonstrated that CD4+ T cells differentiation was unbalanced in PE^[13,16]^. The significant elevation of CD4+ T cells suggests a potential shift in the immune response, which may contribute to the disease’s progression. For instance, the expansion of CD4+ T cells has been linked to increased inflammatory responses and altered immune regulation, which are characteristic features of PE^[32,47]^. This finding aligns with previous studies that an imbalance in immune responses, particularly involving T helper cell subsets, contributes to the inflammatory milieu observed in PE^[48,49]^. Our single-cell RNA sequencing analysis revealed a complex cellular environment in PE, with a notable increase in the proportion of CD4+ T cells, which were found to be enriched in inflammatory cytokines. And immune cells and cytokines are recognized to have a significant function in reshaping the vasculature in microenvironments^[50–52]^,which might contribute to compromised angiogenesis and the onset of preeclampsia and other placental insufficiency disorders ^[53,54]^. This suggests these immune cells may exacerbate the pathological processes associated with PE by promoting vascular inflammation and endothelial dysfunction ^[55]^. Moreover, prior reports have demonstrated that immunoregulatory cytokines and angiogenic factors at the fetal-maternal interface could result in inadequate trophoblast differentiation migration and invasion^[45,56]^, which are also critical features of the disease^[57]^. Furthermore alterations in immune cell populations, along with cytokine generation by CD4+cells, pointed to a weakened response with an exhausted phenotype involving reduced IL1b^[12]^. Therefore, The identification of CD4+ T cells as key players in this context underscores their potential as therapeutic targets. By modulating the activity or polarization of these cells, it may be possible to mitigate the inflammatory responses that lead to adverse pregnancy outcomes. Furthermore, the correlation between the elevated levels of IL-1 cytokines and PE indicates that these cytokines could serve as biomarkers for disease progression, providing a basis for early diagnosis and intervention strategies ^[47]^. Overall, this research not only elucidates the immunological underpinnings of PE but also opens avenues for novel therapeutic approaches aimed at restoring immune balance and improving maternal-fetal health outcomes.

In this study, we demonstrated a significant decrease in both the total number and intensity of interactions in PE compared to the matched control group. This finding is important as it indicates an overall reduction in intercellular communication during severe PE, possibly resulting in impaired immune responses. The unique signaling patterns, especially the changes in EGF, OSM, and SPP1 signal intensities from CD4+ T cells, emphasize the relevance of these pathways in the progression of PE^[58]^. Moreover, our analysis of receptor-ligand pairs involved in modulating communication between key immune cells reveals that the SPP1 signaling pathway is crucial in the context of PE. Specifically, the interaction between SPP1 from macrophages and its corresponding receptors on CD4+ T cells was significantly higher in the control group but absent in the severe PE group. This finding suggests a compromised SPP1-mediated signaling pathway in severe PE. Overall, our results imply that the disrupted communication through the SPP1 pathway may play a vital role in the dysregulation of CD4+ T cell function in severe preeclampsia, highlighting potential therapeutic targets to restore immune balance during pregnancy ^[59]^.

By applying scRNA-seq, alterations in gene expression can be pinpointed to an individual cell, which can partially clarify the heterogeneous immune response of diseases. Our study identified nine intersecting genes significantly expressed in CD4+ T cells during PE, which may influence placental function and hormonal balance, thereby contributing to the disease’s pathogenesis. Among these genes, LCP1 (Lymphocyte Cytosolic Protein 1) plays a crucial role in cytoskeletal organization and cell migration, which is essential for T cell activation and function. Its upregulation in PE suggests a potential mechanism by which CD4+ T cells may enhance inflammatory responses, impacting placental health. PFKL (Phosphofructokinase, Liver) is involved in glycolysis and energy metabolism, and its altered expression may reflect the metabolic shifts in T cells during PE, indicating a link between energy metabolism and immune response in the context of pregnancy. TSPYL2 (Testis-specific Y-encoded-like protein 2) has been associated with cell proliferation and apoptosis, suggesting that its dysregulation could affect T cell survival and function in PE. SREBF1 (Sterol Regulatory Element Binding Transcription Factor 1) is critical for lipid metabolism and has been implicated in the regulation of inflammatory processes, which may further elucidate the role of lipid homeostasis in the immune response during PE. The findings highlight the importance of these genes in mediating the immune response and their potential as therapeutic targets for managing PE. The expression patterns of these genes underscore the complex interplay between immune regulation and placental function, suggesting that interventions targeting these pathways could mitigate the adverse effects of PE on maternal and fetal health. Understanding the roles of these genes in the context of PE may provide insights into novel diagnostic and therapeutic strategies for this condition, which remains a significant cause of maternal and neonatal morbidity and mortality.

The results of the Gene Ontology (GO) and Kyoto Encyclopedia of Genes and Genomes (KEGG) analyses indicate that the pathways enriched in CD4+ T cells are primarily involved in the negative regulation of insulin secretion and the AMP-activated protein kinase (AMPK) signaling pathway. Insulin secretion is a critical process in maintaining glucose homeostasis, and its dysregulation is often implicated in various metabolic disorders, including PE. The negative regulation of insulin secretion suggests that CD4+ T cells may play a role in modulating the endocrine function of pancreatic beta cells, potentially contributing to the pathophysiology of preeclampsia by influencing maternal metabolic status. The AMPK signaling pathway is known to be a key regulator of cellular energy homeostasis and is activated under conditions of low energy availability. This pathway’s involvement indicates that CD4+ T cells may also influence metabolic processes in peripheral tissues, thereby affecting overall maternal health during pregnancy. The identification of these pathways highlights the potential mechanisms through which CD4+ T cells could impact PE. Understanding the interplay between immune responses and metabolic regulation is crucial for developing interventions aimed at improving outcomes for both mothers and their offspring in the context of PE and related disorders.

Nevertheless, this study recognizes its own limitations. Firstly, the restricted sample size in the current study might introduce bias, and thus, larger cohorts and longitudinal analyses are needed to fully elucidate the role of CD4+ T cells over the course of pregnancy. Secondly, despite excellent in silico performance, we had no external gold standard to verify its performance. Thirdly, due to the character of placental tissue sampling, our study design restricts capacity to establish temporality between exposure and outcome to rule out reverse causation. Future investigations should embrace more extensive clinical sample examinations to verify these findings and expedite their transformation into clinical practices.

In summary, our findings underscore the importance of CD4+ T cells in the context of PE and placental biology, suggesting that they may serve as both biomarkers and therapeutic targets for this condition. The implications of these results extend to future research directions, where understanding the mechanistic pathways involving CD4+ T cells could lead to novel interventions aimed at modulating immune responses in placental disorders encompassing PE. Overall, the identification of these immune cell dynamics not only enhances our understanding of preeclampsia but also opens avenues for targeted therapies that could improve maternal and fetal outcomes ^[28,60]^.

### Ethics approval and consent to participate

The study was approved by the Ethics Committee of Shanghai First Maternity and Infant Hospital of Tongji University (No. 2019-034)

## Acknowledgements

We thank the study participants for permitting us to use their personal data.

## Availability of data and materials

Data and materials supporting the findings of this study are available from the corresponding author upon reasonable request. The other data generated and analyzed during the current study are available in the Gene Expression Omnibus (GEO) repository.

## Author contributions

Interpretation of data, drafting the manuscript: WS; Interpretation of data, drafting the manuscript : ZTF, MYX; Analyzing data, reviewing and editing: CY, ZCC, PJ, LL; Conception and design, interpretation of data, validation: LWJ,GW,LZW; Conception and design, interpretation of data, revision: HXL. All authors have read and approved the final manuscript.

## Consent for publication

Not applicable.

